# Heat dissipation capacity influences reproductive performance in an aerial insectivore

**DOI:** 10.1101/711341

**Authors:** Simon Tapper, Joseph J. Nocera, Gary Burness

**Affiliations:** Environmental and Life Sciences Graduate Program, 1600 West Bank Dr., Peterborough ON, Canada; University of New Brunswick, Forestry and Environmental Management, Fredericton NB, Canada; Department of Biology, Trent University, 1600 West Bank Dr., Peterborough ON, Canada

**Keywords:** Energy expenditure, thermal constraint, parental care, climate, activity

## Abstract

Climatic warming will likely increase the frequency of extreme weather events, which may reduce an individual’s capacity for sustained activity due to thermal limits. We tested whether the risk of overheating may limit parental provisioning of an aerial insectivorous bird in population decline. For many seasonally breeding birds, parents are thought to operate close to an energetic ceiling during the 2-3 week chick-rearing period. The factors determining the ceiling remain unknown, although it may be set by an individual’s capacity to dissipate body heat (the heat dissipation limitation hypothesis). To test this hypothesis, over two breeding seasons we experimentally trimmed the ventral feathers of female tree swallows (*Tachycineta bicolor*) to provide a thermal window. We then monitored maternal provisioning rates, nestling growth rates, and fledging success. We found the effect of our experimental treatment was context-dependent and varied with environmental conditions. Females with trimmed plumage fed their nestlings at higher rates than controls when conditions were hot and non-windy, but the reverse was true under cool and windy conditions. On average, nestlings of trimmed females were heavier than controls, and had a higher probability of fledging. We suggest that removal of a thermal constraint allowed females to increase provisioning rates, but additionally provided nestlings with a thermal advantage via increased heat transfer during maternal brooding. Our data provide partial support for the heat dissipation limitation hypothesis, and suggest that depending on weather patterns, heat dissipation capacity can influence reproductive success in aerial insectivores.

## Introduction

With rising global temperatures, animals will experience warmer than average annual temperatures, and increased frequency of heat waves and droughts (IPCC 2014). Under such changing conditions, animals must maintain stable body temperatures (T_b_) in the face of heat stress (McKechnie and Wolf 2009). A rise in T_b_ above steady state levels (i.e., hyperthermia) occurs when heat is generated and/or acquired from the environment faster than it can be dissipated (Speakman and Król 2010). Non-fatal hyperthermia can have several deleterious physiological consequences, including disrupted cellular signalling (Boulant 1998); impaired synthesis and damage of proteins (Roti Roti 2008); elevated levels of oxidative stress (Costantini et al. 2012); depressed innate and adaptive immune function (Palermo-Neto et al. 2013); and impaired growth and development (Baumgard and Rhoads 2013), among others.

Given the suite of physiological consequences that can develop from hyperthermia, the capacity to dissipate body heat has recently been proposed as a key factor shaping the behaviour, physiology, and ecology of endotherms (“Heat Dissipation Limitation” (HDL) hypothesis (Speakman and Król 2010). Specifically, the HDL hypothesis posits that in endothermic animals maximally sustained energy expenditure is limited by an individual’s maximal capacity to dissipate body heat. Understanding the limits to sustained energy expenditure, or sustained metabolic rate (SusMR), is important because metabolic ceilings could impose constraints on life-history traits (Drent and Daan 2002; Peterson et al. 1990). For example, an energetic ceiling in chick-rearing birds could theoretically influence clutch size because parent birds can only feed a certain number of chicks based on their sustained level of energy expenditure (Peterson et al., 1990; Monaghan and Nager, 1997).

There are several lines of indirect evidence, which support the HDL hypothesis. For instance, activity levels decline with high ambient temperature (T_a_) (Carroll et al. 2015; Theuerkauf et al. 2003; Zub et al. 2013); animals preferentially select habitats within their home range to escape solar radiation at the expense of resource acquisition (Lease et al. 2014; Street et al. 2015; van Beest et al. 2012); in birds, individuals with bigger bills (larger “thermal windows”) are more active and spend more time singing on hot days than do birds with smaller bills (Luther and Danner 2016).

While there exists indirect evidence from studies across different taxa, direct tests of the HDL hypothesis have primarily been performed on lactating laboratory rodents. For example, when lactating animals are experimentally exposed to cooler temperatures, they can increase energy intake and milk production beyond levels seen at warmer ambient temperatures (Hammond et al. 1994; Johnson and Speakman 2001; Ohrnberger et al. 2016). Furthermore, when the fur of lactating rodents is shaved, they consume more food and produce more milk compared with non-shaved controls (Gamo et al. 2016; Król et al. 2007), suggesting that capacity to dissipate heat place limits on performance.

Despite this direct evidence of an HDL for laboratory mammals, there have been few experimental studies testing the HDL hypothesis in free-ranging animals (Valencak et al., 2011;). Two recent studies on breeding tits, however, provided some experimental evidence for the HDL hypothesis in free-ranging birds (Nilsson and Nord 2018; Nord and Nilsson 2018). Nord and Nilsson (2018) removed the ventral plumage from breeding blue tits, *Cyanistes caeruleus*, and found that feather-clipped parents had heavier nestlings than controls, while maintaining lower body temperatures and without additional mass loss. However, there were no differences in feeding rates between the treatments, and only older parents had heavier nestlings. This result suggests that the limit to SusMR might be influenced by life-history strategy, which could also explain why some species (e.g., European Hare, *Lepus europaeus*, Pallus) do not exhibit an HDL (Valencak et al. 2011).

One aspect largely ignored by previous studies examining the limits to SusMR is the influence of other environmental factors on heat balance. For instance, wind carries heat away from the body via convection, and increasing wind speeds decrease heat gain from solar radiation (Wolf and Walsberg 1996). Further, rates of evaporative water loss can be severely inhibited by high humidity, and thus variation in humidity could alter activity levels (Gerson et al. 2014). Finally, in addition to the temperature, wind speed, relative humidity, and precipitation have been shown to play an important role in influencing foraging activity in aerial insectivores (Cox et al. 2019; Ouyang et al. 2015; Rose 2009). Therefore, testing the HDL hypothesis in free-ranging animals should attempt to control for these additional factors.

To determine whether heat dissipation constrains reproductive performance in breeding birds, we experimentally manipulated the ability of female tree swallows (*Tachycineta bicolor*) to dissipate body heat, by removing feather overlying the brood patch. Tree swallows are an excellent model species in which to test the HDL hypothesis. As aerial insectivores, they are active foragers, and can spend up to16 hours per day gathering insects to feed their nestlings. We predicted that, if in general (i.e., across environmental conditions), the ability to dissipate body heat limits SusMR then (1) trimmed birds would maintain higher feeding rates than control birds (2) trimmed birds would have heavier offspring and (3) offspring of trimmed birds would have greater fledging success than those of non-trimmed birds. Given that temperature does not occur in isolation of other environmental factors, these predictions maybe context-dependent.

## Materials and methods

### Study area and species

All research was approved by the Trent University Animal Care Committee, in accordance with the Canadian Council on Animal Care (AUP # 24747). We conducted this study in May-July 2017 and 2018, on two nest-box breeding populations of tree swallows located at the Trent University Nature Areas, Peterborough, Ontario, Canada (44°21′N, 78°17′W) and at the Lakefield Sewage Lagoon, Lakefield, Ontario (44°24’58.3”N 78°15’26.8”W). The Trent Nature Areas consists of relatively open grassy fields, and there are about 70 boxes spaced ∼10-20m apart. The Lakefield Sewage Lagoon consist of two rectangular lagoons, with 50 boxes encircling the perimeter, and spaced 10-20m apart. Females at both sites typically lay clutches of five to seven eggs, with one egg laid each day. Once a clutch is completed, females incubate the nest for approximately 14 days, and nestlings typically hatch synchronously. Nestlings typically fledge 18-22 days post-hatch.

### General field methods

Beginning in May each year, we checked nest boxes every other day until the presence of nest material was discovered, at which point boxes were monitored every day until clutch completion. We used a marker pen to sequentially number eggs as they were laid; the date the last egg was laid was considered to be day 0 of incubation. Hatch date (day 0) for the brood was considered the first day when nestlings hatched.

### Experimental manipulation

We captured females during early nestling provisioning (see capture protocol below), and upon capture, randomly assigned females to either a trimmed or control condition, based on a flip of the coin. In the trimmed condition, we removed the contour and downy feathers covering the brood patch (details below) to expose the bare skin underneath (Figure S1). We chose to remove feathers from this region because 1) it is highly vascularized, increasing the chance of heat loss and 2) there would be minimal interference with flight.

We performed trimming manipulations with two people: one person held the bird ventral side up, while the other person, using surgical scissors, cut the feathers away. Control females were handled identically, but instead of cutting the feathers, we performed a “mock cut”, in which we cut the air above the brood patch. In 2018, we additionally measured the size of the exposed area for all trimmed females (n = 21). We quantified both the length and width of the exposed skin using a piece of string. The median (± median absolute deviation, MAD) length of exposed skin was 3.1 ± 0.2cm (range: 2.6-3.7cm) and the median width of exposed skin (± MAD) was 1.9 ± 0.1cm (range: and 1.5-2.1cm). Assuming the trimmed area was an ellipse, the amount of exposed skin would be 4.63cm^2^. The estimated percentage of total surface area trimmed was ∼7% (see supplementary material for details).

### Remote monitoring of activity

As an index of activity, we quantified provisioning rate of females using passive integrative transponder (PIT) tags. During late incubation (day 7 – day 10 post-clutch completion), we captured females in their nest box and implanted them in the nape of the neck with either 1) non-temperature sensitive (EM4100; #11001, GAO RFID, Ontario, Canada) or 2) temperature-sensitive (Biotherm13; Biomark, Boise, Idaho, USA) PIT tags, following Nicolaus et al. (2011). Data on body temperature from the Biotherm13 tags are part of a parallel study. Following implantation, we recorded body mass, wing chord (flattened), head-bill length, exposed culmen (*sensu* Borras et al., 2014), and determined age (second year or after second year) based on plumage coloration (Hussell 1983). Total time in the hand was approximately 12 minutes. Details regarding the reader set-up are described in the supplementary materials.

On day 1 post-hatch, we captured females again and performed the experimental manipulation (control vs. trimmed), recorded body mass, and obtained a 50-75 μL blood sample from the brachial vein as part of a parallel study. At day 10 of provisioning, we again measured body mass and collected a second blood sample.

We had six females with data from both years. We attempted to give each bird the opposite treatment that it received in 2017, but in an effort to keep the sample sizes within the treatments approximately balanced, four individuals received the opposite treatment and two received the same treatment.

### Nestling measurements

Nestlings were measured between ∼1200 – 1800h. To determine the effect of maternal treatment on nestling growth rate, we weighed nestlings at days 0 (hatch), 3, 6, 9 and 12 (i.e., peak body mass). We did not handle nestlings beyond day 12 to prevent premature fledging. In 2017, we weighed nestlings on a spring and digital scale and in 2018 on a digital scale only. This did not affect our conclusions, but see supplementary material for details. As an index of nestling body size, we measured wing chord on day 12 using a wing-rule (with a stop). As part of another study, we collected a blood sample (∼75 μL) from each nestling at day 12. Fledging success was determined after checking all nest boxes on day 18 post-hatch, and in the following days as necessary. There were no instances of premature fledging from checking the nest box.

### Data compilation and organization

#### Feeding rate

Most adult females were caught when nestlings were 1-2 days of age (see *female captures* above), and we therefore only included feeding rate data between nestling ages 3 to 14d, and between the hours of 0500 and 2100. We only included data collected within 0500 and 2100h because swallows are relatively inactive during this window (S.Tapper, unpublished data). Our feeding rate data range from 01-Jun-2017 to 29-Jun-2017 and from 31-May-2018 to 11-Jul-2018.

For data organization and statistical analyses, we used R (version 3.5.1, R Core Team). To transform raw RFID reads into visits to the nest box, we used the function “visits” from the package feedr (LaZerte et al. 2017). We considered repeated reads from the same individual as a singular event if successive reads were < 60s apart. We defined daily feeding rate as the total number of visits per day / the total number of hours in which the bird was logged on the reader (i.e., an integer). We included each hour toward the total number of hours spent provisioning if there was at least one read in the hour of interest. We chose this definition over a typical standardized provisioning rate (e.g., number of visits/16hrs (0500-2100h) because of unequal number of observations across birds. Unequal observations were due to 1) birds with thermal tags having fewer overall hours of data (due to cycling of readers among boxes) and 2) because some birds had missing data as a result of equipment failure. This definition provides a relatively unbiased measure of feeding rate compared with one in which the total number of hours across all birds was assumed to be the same (i.e., standardized).

#### Environmental variables

We gathered data on daily mean ambient temperature (°C), wind speed (km·hr^−1^), relative humidity (%), and total precipitation (mm), from Trent University’s weather station, which is located approximately 1.5km from the Trent University Nature Areas and 9.5km from the Sewage Lagoon and (downloadable from Environment Canada, http://climate.weather.gc.ca/index_e.html). We calculated averages over 24h, because overnight and early morning weather could have carry-over effects on feeding behaviour the next day. To avoid multiple univariate statistical tests we collapsed our four weather variables using principle component analysis (PCA). We centered and scaled the data prior to calculation of the correlation matrix. The first two PCs explained a combined total of 69.1% (PC1 = 41.54%, PC2 = 27.58%) of the variation in weather. PC1 was loaded primarily by relative humidity and total precipitation (Table 1), while PC2 was predominately loaded by wind speed and temperature (Table 1). We included both PC1 and PC2 scores in our statistical models because we were interested in how the different weather variables related to treatment independently of each other (i.e., humidity and precipitation vs. wind speed and temperature),

**Table 1.**
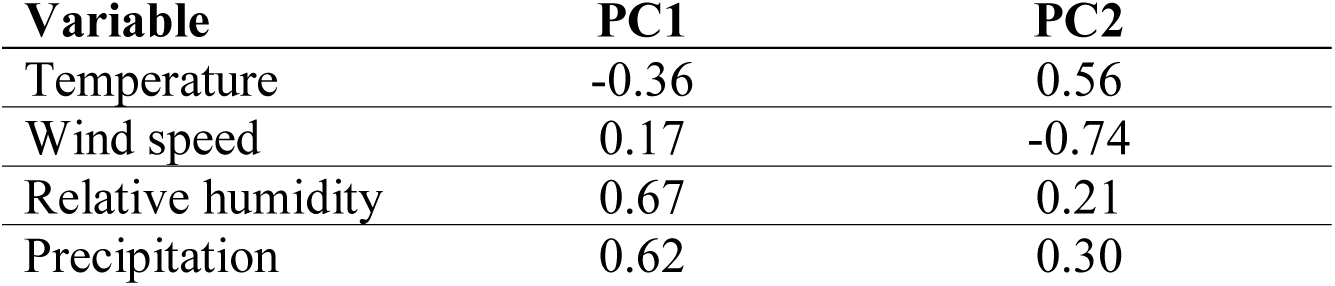
Rotated matrix values of principal component analysis scores for 2017 and 2018 environmental data. To improve interpretation of results, we multiplied PC1 by −1 so that relative humidity and precipitation were positively related to PC1 score. The first two PCs explained a combined 69.1 % variance in the weather data (PC1 = 41.54 %, PC2 = 27.58 %).

| **Variable** | **PC1** | **PC2** |
| --- | --- | --- |
| Temperature | -0.36 | 0.56 |
| Wind speed | 0.17 | -0.74 |
| Relative humidity | 0.67 | 0.21 |
| Precipitation | 0.62 | 0.30 |

### Statistical analyses

For all analyses (i.e., feeding rate, nestling body mass, fledging success), we only included nests that fledged at least one nestling. Unless otherwise stated, model parameters were estimated by restricted maximum likelihood, degrees of freedom and *p*-values were calculated using the Sattherwaite approximation in the lmerTest package (version 3.1-0, Kuznetsova et al., 2017), and confidence intervals were calculated with the Wald method in the lme4 package (version 1.1-20, Bates et al., 2015). Means reported are estimated marginal means, generated using the emmeans package (version 1.3.2, Lenth, 2019). We considered *P*-values ≤ 0.05 as statistically significant. We checked that our models met assumptions of normality and homogeneity of variance by visual inspection of the residuals.

### Feeding rate

We tested for an effect of experimental treatment (trimmed or control) on maternal feeding rate (expressed as visits·hr^−1^) using linear mixed effects models (lme4). In 2017, there were a total of 18 females included in the analysis (Control_2017_ = 10, Trimmed_2017_ = 8), while in 2018 there were a total of 37 females (Control_2018_ = 18, Trimmed_2018_ = 19). We included brood size, treatment, PC1 score, PC2 score, year, lay date, and chick age as main effects in our model, and interactions terms for: (1) treatment × PC1, and (2) treatment × PC2. In each year, we standardized lay date so that the first day in which a female laid an egg was given the value of 0. We controlled for repeated observations from the same individual across days by including bird identity as a random effect.

After running the initial model and assessing diagnostic plots, we detected one observation with a standardized residual that was > 3 standard deviations above 0 (all others < 2.8), which we considered an outlier. Exclusion of this point did not change our results, but improved model fit, and so was excluded from the analysis.

### Nestling morphology and fledging success

We tested for differences in nestling growth rates between treatments using a three-parameter logistic growth curve, which has been shown to model tree swallow growth accurately (McCarty 2001; Zach and Mayoh 1982). Our sample size in 2017 and 2018 was 19 (Control_2017_ = 12, Trimmed_2017_ = 9) and 37 (Control_2018_ = 18, Trimmed_2018_ = 19) nests respectively. At each time point on the growth curve (i.e., 0, 3, 6, 9, 12d post-hatch), we calculated the average nestling mass, per brood, and used this as our dependent variable, because we did not track individual nestlings in 2017.

We constructed the growth curve model using the “nlme” function from the nlme package (version 3.1-137, Pinheiro et al., 2014). To describe the patterns of nestling growth, we calculated three parameters from the growth curve: the asymptotic mass (*A*) (i.e., peak mass, ∼ day 12 post-hatch) (in grams), the inflection point (i.e., point of steepest growth) (*I*) of the growth curve (in days), and the growth rate constant (i.e., steepness of growth curve) (*K*). We estimated our parameter starting values using the “SSlogis” function from Stats package (base R).

We included ‘maternal identity’ as a random intercept on the asymptotic parameter to control for statistical non-independence in the growth rate among nestlings that were dammed from the same females. A random intercept for ‘maternal identity’ was initially applied to all growth rate parameters (*A, K, I*), however, application of a random intercept to the asymptotic parameter alone explained the greater variance in our data (see supplementary material, Table S1, for more details). Confidence intervals and predictions were calculated using bootstrapping with replacement based on 1000 replications.

We tested for differences in day 12 nestling wing length between treatments using a linear mixed effects model (lme4). Our model included main effects of treatment, lay date (standardized), and year, and maternal identity was treated as a random effect to control for both statistical non-independence of returning mothers (n=6) between years and nestlings within the same brood.

To determine whether treatment affected an individual nestling’s fledging success, which we defined as either 1 (fledged) or 0 (did not fledge), we used a generalized linear mixed model (glmer function in lme4) with a binomial error distribution and a logit link. We used the same model structure as we did for nestling morphology. After plotting the predictions from the model, we noticed differences in variance between treatments, and subsequently ran an F-test (using the var.test function in stats package, base R) on the predicted probabilities from the model. Results from the F-test confirmed violation of homogeneity of variance (F_163,145_ = 6.78, 95% CI [4.93, 9.31], *P* < 0.0001) and we re-ran our model with the inclusion of a variance structure (using with the “weights” argument) to control for heteroskedasticity between treatments.

## Results

### Feeding rate

On average, maternal feeding rate (visits·hr^−1^) (± SE) did not differ between treatments (*P* = 0.165; control birds: 11.8 ± 0.55, trimmed birds: 11.4 ± 0.53). Feeding rate was negatively related to PC1 score (*P* < 0.0001, Fig. 1, Table 2), indicating that birds foraged less on wetter and more humid days. We did not detect an interaction between treatment and PC1 score (*P* = 0.122, Table 2), but there was a trend of higher feeding rates in control birds on wetter and more humid days relative to trimmed birds (Fig. 1). Feeding rate differed significantly between treatments as a function of PC2 score (i.e., treatment × PC2 score, *P* = 0.020, Fig. 2, Table 2). At the highest PC2 score (i.e., 1, indicating high temperature, low wind speed), trimmed birds made 5.5% more trips per hour (∼8 extra visits, given a 16hr day) than controls. However, at the lowest PC2 score (i.e., 0, meaning low temperature, high wind speed), trimmed birds made 23% less trips per hour (∼32 visits, given a 16hr day) than control birds. Feeding rate increased with brood size (*P* = 0.045, Table 2); females raising larger broods (7 nestlings) made ∼3 more visits to the nest per hour than mothers raising small broods (3 nestlings). Provisioning rate was also negatively related to lay date (*P* = 0.055, Table 2), meaning that earlier nesting birds had higher feeding rates than later nesting birds.

**Table 2.** Factors contributing to variation in maternal feeding rate. Fixed effect coefficient estimates with 95% confidence intervals (CI), and *P*-values.

| **Predictor** | **Estimate** | **95% CI** | ***P*-value** |
| --- | --- | --- | --- |
| Intercept | 10.13 | 5.87 – 14.39 | <0.001 |
| Treatment | -1.30 | -3.13 – 0.53 | 0.165 |
| PC1 | -5.48 | **-7.12 – -3.84** | **<0.001** |
| PC2 | 0.75 | -0.98 – 2.49 | 0.394 |
| Year | 0.01 | -0.97 – 0.99 | 0.977 |
| Lay date | -0.12 | -0.24 – 0.00 | 0.055 |
| Brood size | 0.74 | 0.03 – 1.45 | 0.045 |
| Nestling age | 0.01 | -0.06 – 0.07 | 0.856 |
| Treatment x PC1 | -1.86 | -4.22 – 0.50 | 0.123 |
| Treatment x PC2 | 2.84 | **0.46 – 5.22** | **0.020** |

**Fig. 1.**
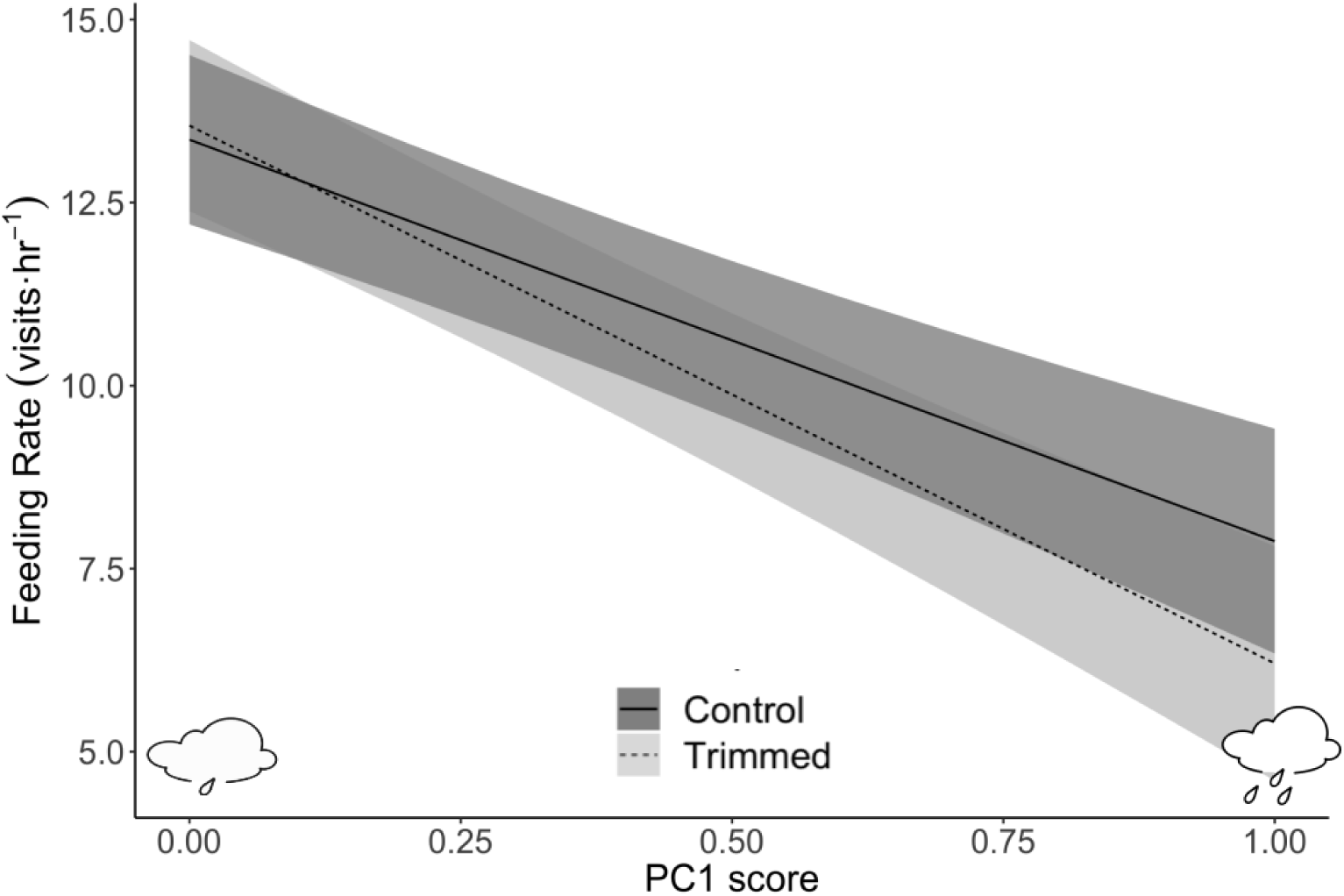
Prediction plot of female provisioning rate (± 95% CI) in relation to PC1 score. All other predictors in the model are held at their constant value. Low PC1 scores represent low rainfall and low humidity conditions, while high PC1 scores represent high rainfall and high humidity conditions. As relative humidity and rainfall increased, females with an increased capacity to dissipate heat (i.e., trimmed) decreased feeding rates more sharply than control birds. Sample sizes (N_control_= 28, N_trimmed_=27).

**Fig. 2.**
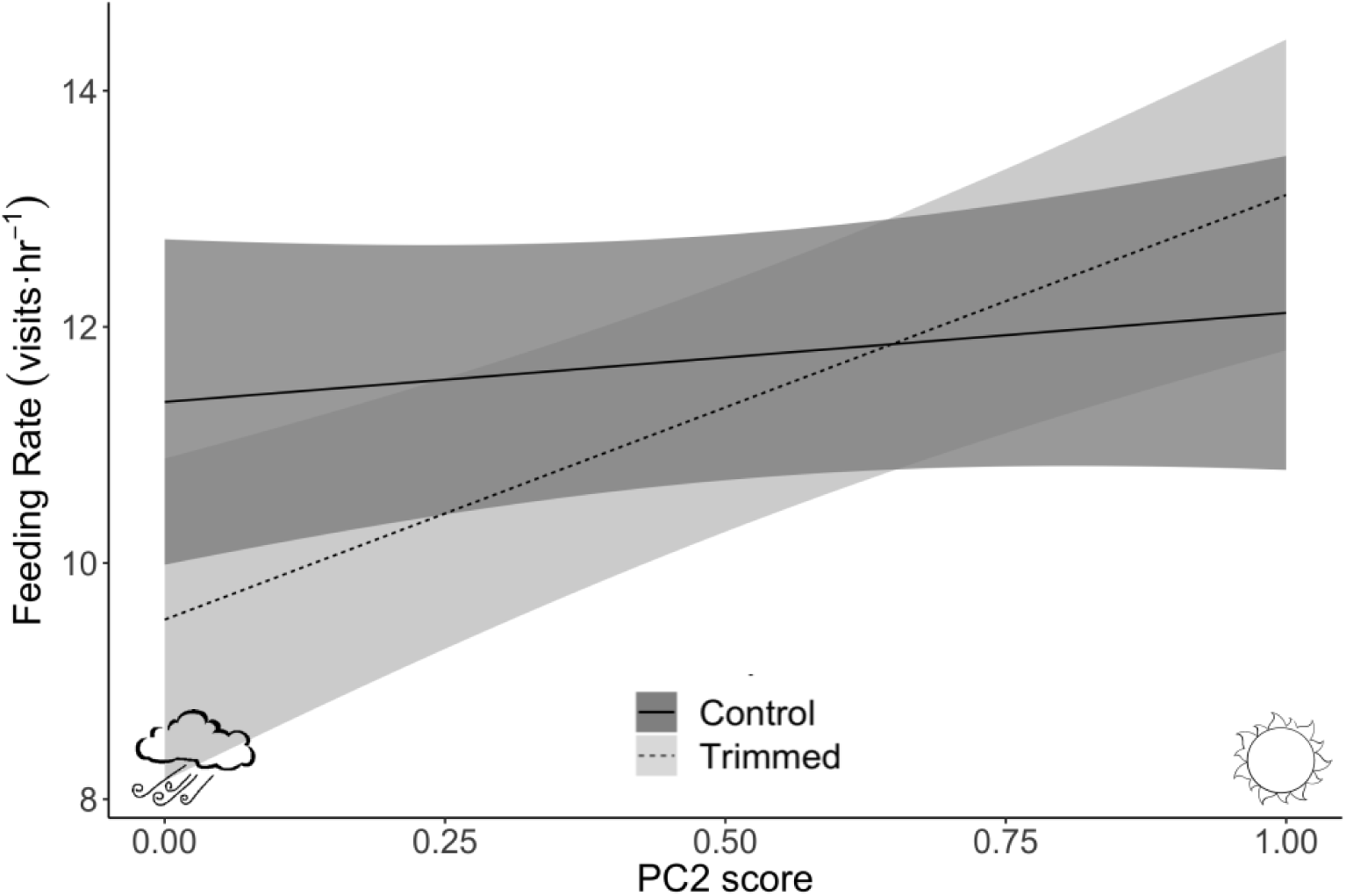
Prediction plot of female feeding rate (± 95% CI) in relation to PC2 score. All other predictors in the model are held at their constant value. Low PC2 scores represent cool and windy conditions, while high PC2 scores represent hot and calm conditions. As temperature increased and wind speed decreased, females with an increased capacity to dissipate heat (i.e., trimmed) increased feeding rates, while control birds remained relatively constant across PC2 scores. Sample sizes (N_control_= 28, N_trimmed_=27).

### Nestling morphology and fledging success

The best model for describing nestling mass included a random intercept on asymptotic mass (supplementary material, Table S1). Nestlings from trimmed mothers were heavier (± SE) by 1.71 ± 0.48g at their asymptote (∼ day 12 post-hatch) compared with nestlings from control mothers at their asymptote (i.e., Treatment, *P* = 0.001, Fig. 3, Table 3). We did not detect any significant differences in the inflection point (∼ day 5 post-hatch) between groups (*P* = 0.061, Table 3), nor in the growth rate constant (i.e., steepness of curves) between groups (*P* = 0.548). Wing length did not statistically differ between treatments (*β* = 0.18, 95% CI [-0.03, 0.38], *P* = 0.090), although nestlings in 2017 had longer wings than nestlings in 2018 (i.e., Year, *β* = −0.38, 95% CI [−0.57, −0.20], *P* < 0.001). Lay date was negatively related to wing length (*β* = −0.03, 95% CI [−0.05, −0.01], *P* = 0.006).

**Table 3.** Parameter estimates for nestling growth trajectories. Fixed effect coefficients with 95% confidence intervals (CI), and *P*-values.

| **Parameter** | **Predictors** | **Estimates** | **95% CI** | ***P*-value** |
| --- | --- | --- | --- | --- |
| Asymptote<br>(*A*) | Intercept | 21.1 | 20.15 – 21.67 | <0.001 |
|  | Treatment | 1.68 | **0.37 – 2.72** | **0.001** |
| Inflection point<br>(*I*) | Intercept | 4.75 | 4.65 – 5.08 | <0.001 |
|  | Treatment | 0.21 | -0.08 – 0.51 | 0.064 |
| Growth rate constant<br>(*K*) | Intercept | 1.91 | 1.76 – 2.01 | <0.001 |
|  | Treatment | 0.05 | -0.12 – 0.23 | 0.548 |

**Fig. 3.**
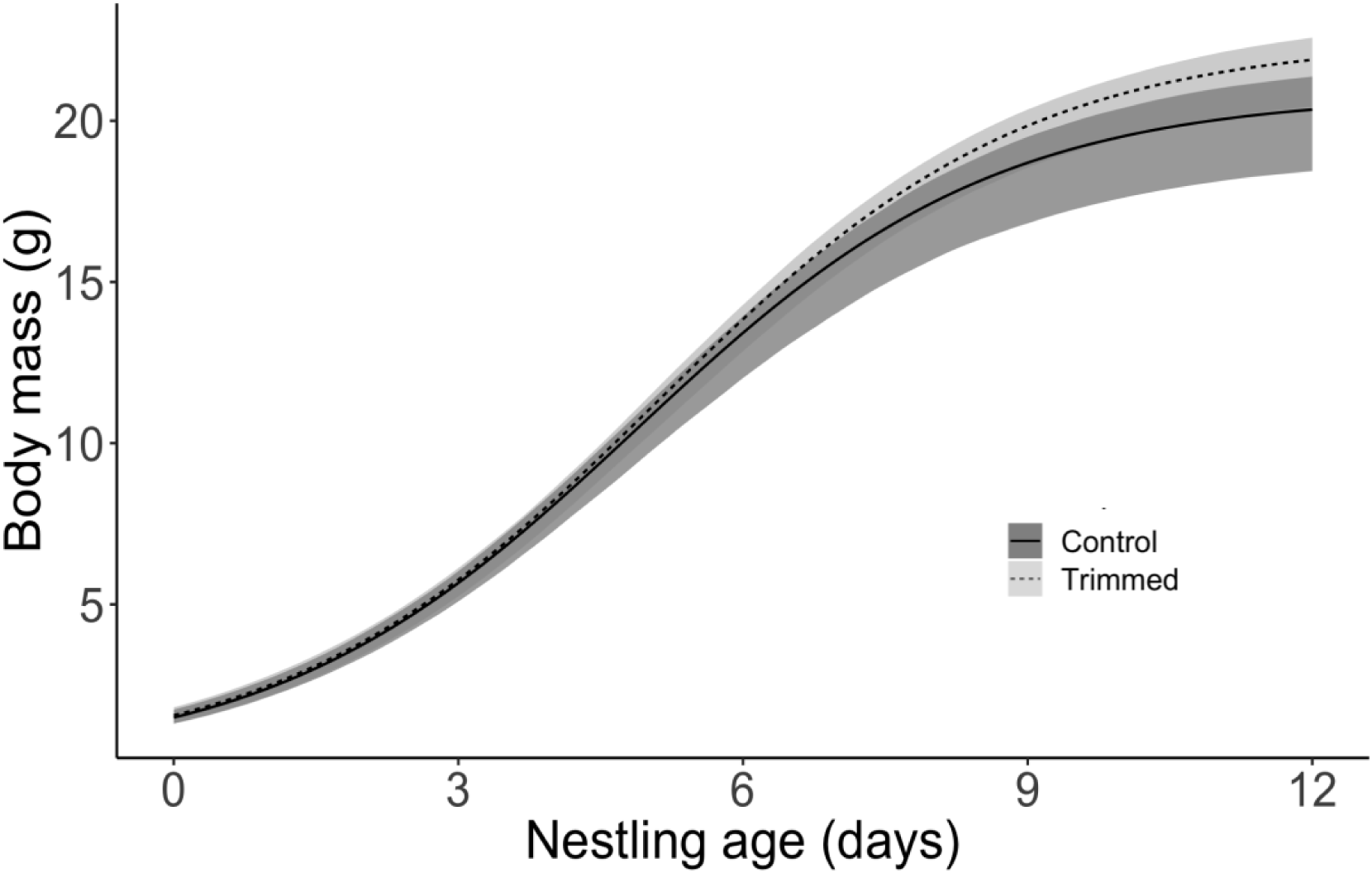
Estimated growth trajectories comparing the growth of nestlings from control broods and from those in which females had increased heat dissipation capacity (i.e., trimmed). Bands around lines represent the 95% confidence intervals obtained from bootstrapping. There were 309 nestlings from 58 nests (N_Control_=30, N_Trimmed_=28).

The probability of fledging (± SE) was significantly higher for nestlings in trimmed (98.4 ± 0.03%) compared to control broods (94.1 ± 0.01%) (Odds Ratio = 3.93, 95% CI [1.03, 14.96], *P* = 0.045, Fig. 4), and did not differ significantly between years (Odds Ratio = 2.62, 95% CI [0.77, 8.89], *P* = 0.122). Similar to nestling morphology, lay date was negatively related to fledging success (Odds Ratio = 0.87, 95% CI [0.76, 1.00], *P* = 0.054).

**Fig. 4.**
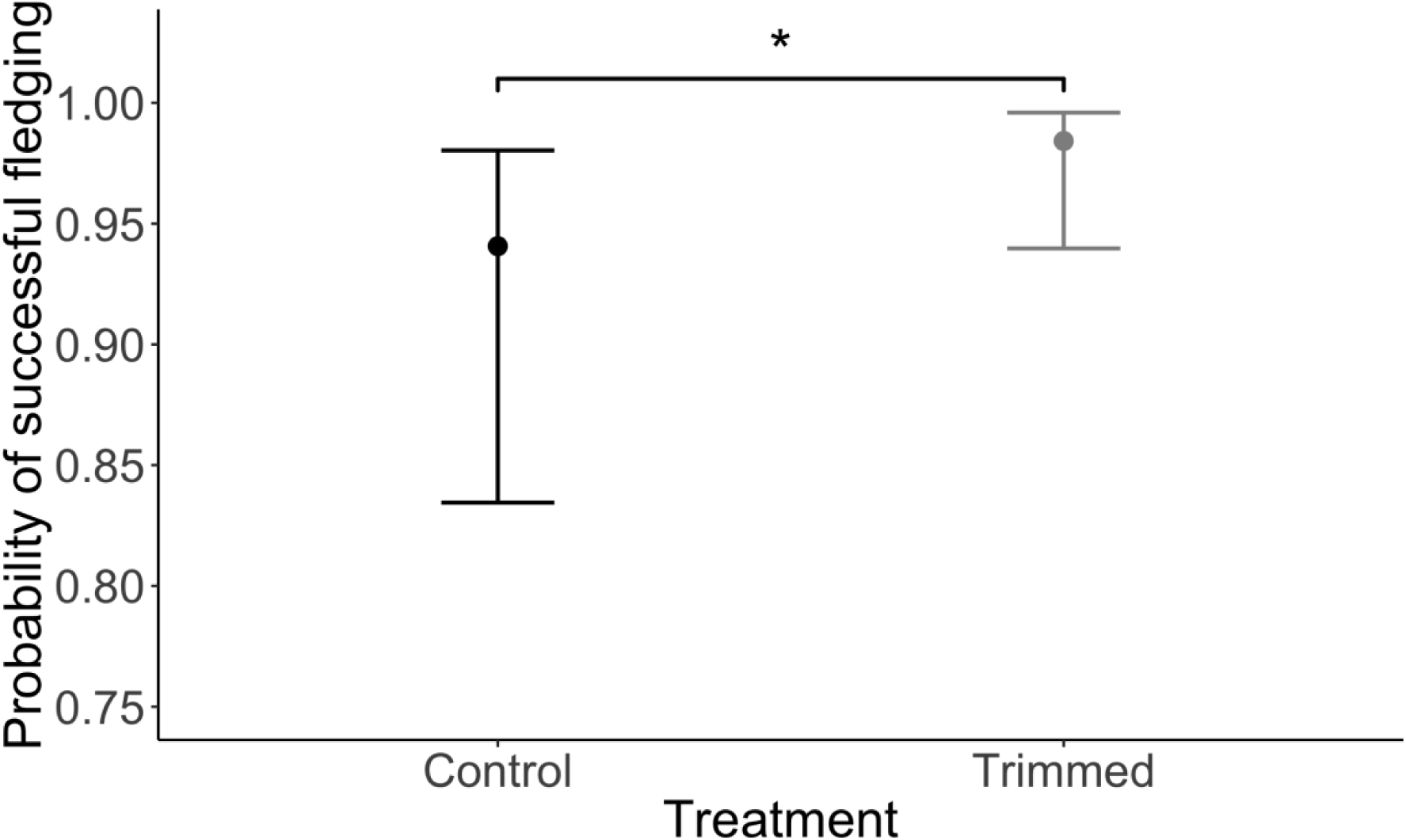
Predicted probability of nestling fledging success as a function of treatment (± 95% confidence intervals), converted from odds ratios to probabilities. * *P* < 0.05. There was a higher probability that nestlings from trimmed broods would fledge over nestlings from control broods. There were 309 nestlings from 58 nests (N_Control_=30, N_Trimmed_=28)

## Discussion

We found that the ability to dissipate body heat affected the reproductive performance of female tree swallows in a context-dependent manner, providing partial support for the HDL hypothesis. We had predicted that trimmed birds would maintain higher feeding rates than control birds due to the increased capacity to dissipate body heat. Instead, an individual’s feeding rate depended on the interactive effects of multiple environmental factors. When it was hot and wind speeds were low (high PC2 score), trimmed birds tended to provision their offspring more than control birds, but this pattern was reversed when it was cool and windy (low PC2 score) (Fig. 2).

The relatively higher activity rates of trimmed birds at high PC2 scores (hot and calm) compared with control birds is consistent with studies from the mammalian literature, in which experimental fur removal in mice allows for higher sustained energy expenditure during lactation, as measured via increases in food intake and milk production (Gamo et al. 2016; Król et al. 2007). It is also consistent with the idea that different mechanisms (e.g., heat dissipation vs. peripheral limitation) can constrain SusMR in different circumstances (Wen et al. 2017). For instance, lactating laboratory mice appear to face a heat dissipation limitation at warm (30 °C), but not room (i.e., 21 °C), temperatures (Wen et al. 2017). Given that the relative effect of PC2 on feeding rate was much weaker at the high end of the scale (i.e., when warm and calm trimmed birds made 5% more trips than controls) versus the low end of the scale (i.e., when cool and windy trimmed birds made 23% fewer trips than controls), our study provides only partial support to the HDL hypothesis.

Wind speed can reduce foraging rates in tree swallows, and daily average wind speed can have a greater effect on feeding rate than average temperature, presumably because high winds make aerial insects more difficult to find and/or by increasing the flight costs during foraging (Rose 2009). If flight costs were increased in windy conditions, thereby increasing energy expenditure and T_b_ (Wolf et al. 2000), trimmed birds would have an advantage because of increased heat dissipation capacity. However, this would only be advantageous if temperatures were also warm, as this would minimize heat gain from solar radiation (Wolf and Walsberg 1996). In the context of our study, temperature and wind were negatively related in PC2, so the birds were not experiencing warm and windy conditions, but rather were experiencing cool and windy conditions. Thus, it is likely that trimmed birds would have difficulty maintaining heat balance in cool air temperatures and high wind speeds, because heat loss to the environment would occur more quickly than any heat generated via metabolism (Zerba et al. 1999).

There was a significant decline in feeding rate with increasing PC1 score (relative humidity and precipitation). While we did not detect a statistically significant interactive effect of treatment and PC1 (*P* = 0.12) on feeding rate, trimmed birds appeared to feed their nestlings less frequently at high PC1 scores (high precipitation, high relative humidity) than at low PC1 scores (low precipitation, low relative humidity) (Fig. 1). Precipitation has been found to negatively influence feeding rate in passerines (Öberg et al. 2014; Winkler et al. 2013), and it is generally assumed that the decline in foraging rate is caused by a reduction in insect activity (Cox et al. 2019; Irons et al. 2017; Winkler et al. 2013). If however precipitation did decrease tree swallow foraging rate due to lowered insect activity, this should have affected both treatments equally, but this tended not to be the case as Fig. 2 shows. Rain has a “cooling effect” on the subjective temperature an animal experiences, which would make thermoregulation more difficult for an animal with less insulation (i.e., a trimmed bird). Furthermore, precipitation is typically associated with cloud cover, which would limit thermal radiation reaching the birds. Therefore, precipitation may have influenced foraging rate because of the challenges of thermoregulating under cool and wet conditions, rather than because of reduced insect activity.

Relative humidity can also influence activity level in birds (Gerson et al. 2014). When T_a_ exceeds T_b_, heat cannot be lost via dry heat transfer, and evaporative heat transfer is the only means to dissipate metabolic heat. While T_a_ never exceeded T_b_ in our study (max T_a_=34°C, mean T_b_=∼ 41°C, S. Tapper, unpublished data), relative humidity still plays an important role in thermoregulation because it decreases the potential for evaporative heat loss (Gerson et al. 2014). In our study, as precipitation and relative humidity increased together in PC1, temperature simultaneously decreased in PC1. Birds in our study were therefore likely experiencing cool and wet conditions, rather than warm and wet conditions. If increased precipitation had a cooling effect on air temperature, then this would explain why trimmed birds had lower foraging rates than control birds at higher PC1 scores.

We predicted that due to increased activity rates, trimmed females would have heavier nestlings than controls. In line with this prediction, trimmed females had nestlings that reached higher asymptotic masses (∼ day 12 post-hatch, Fig. 4) compared with control females. However, the mechanism by which this prediction supports the HDL hypothesis is unlikely to be due to feeding rate alone, because trimmed birds had a relatively minor advantage in feeding rate at high PC1 (hot and non-windy) scores relative to their disadvantage at low PC1 scores (cool and windy). The heavier offspring of trimmed females may in part be due to trimmed females transferring more heat to their offspring during brooding compared with control mothers. Tree swallow nestlings do not develop feathers until around 6 to 7 days post-hatch (Marsh 1980), and mothers typically continue brooding until swallows reach 5 days of age (McCarty 1996). Although we did not estimate heat transfer of the brood patch/trimmed area between treatments, in a study similar to ours, Nord and Nilsson (2018) trimmed ∼22% of the total surface area of male and female blue tits resulting in an approximate 47% increase in estimated heat transfer for trimmed compared to control birds. However, Nord and Nilsson (2018) trimmed a greater proportion of the plumage than we did (22% in their study, 7% in ours), as we did not remove feathers covering the pectoral muscles. Nevertheless, it is plausible that a 7% difference in estimated heat transfer between treatments could lead to an advantage in terms of increased growth and survivorship (Dawson et al. 2005; Klaassen et al. 1994; Pérez et al. 2008). For instance, if nestlings were experiencing cool temperatures and responded by increasing metabolic rate (Nord and Nilsson 2011), then nestlings from trimmed broods receiving direct heat transfer from the enlarged bare skin surface could have had more energy to put into growth rather than maintaining homeothermy.

Nestlings from trimmed broods may also have been heavier than controls because of adults providing nestlings with different quantities, or quality of food (Sofaer et al. 2018; Twining et al. 2016), or because male swallows adjusted their provisioning strategy in accordance with female behaviour, thus leading to differences in the total feeding frequency per nest (Akçay et al. 2016; Lendvai et al. 2018).

We predicted that in addition to producing heavier and structurally larger nestlings at day 12, nestling fledging success would be greater in trimmed compared to control broods. In line with this prediction, fledging success was higher for nestlings from trimmed broods (*P* = 0.045) and was also less variable compared to control broods (Fig. 4). This suggests lower overall mortality for nestlings in trimmed compared to control broods. In birds, fledging success is typically correlated with post-fledging survival and recruitment (McCarty 1996; Weatherhead and Dufour 2000). While we do not have the data to examine post-fledging survival, it is possible that increased fledging success and less variance around fledging success for trimmed birds could also mean less variability in post-fledging survival, which would suggest a possible fitness benefit for trimmed birds.

In conclusion, our data demonstrate that heat dissipation capacity is an important factor influencing female tree swallow behaviour and breeding success. We provide evidence that even a small adjustment to the ventral plumage can cause birds to modulate their activity levels, as measured indirectly via provisioning rate. Therefore, this supports the HDL hypothesis’s predictions that heat dissipation can alter SusMR in free-living animals.

As global temperatures and frequencies of heat waves increase (IPCC 2014), the physiological parameter of heat balance will be of higher concern for all birds, particularly aerial insectivores such as tree swallows. The birds in our study did not experience temperatures beyond their T_b_, but we provide evidence that under such extreme conditions, being unable to effectively dissipate body heat could decrease reproductive success. Our study also highlights the likely reason why birds mostly keep their plumage during the breeding season: because cool, rainy, and windy weather can negatively affect foraging rate and potentially thermoregulation. In addition to warming temperatures, climate change may also reduce the food supply of tree swallows (Irons et al. 2017; Winkler et al. 2013). Tree swallows may then have to contend with both the effects of reduced food supply and overheating while feeding their young.

## Acknowledgements

We thank A. Schubert and J. Baici for help with data collection, D. Marshall for help with antenna construction, E. Bridge for assistance with data loggers, A. Gerson for loaning RFID equipment, J. Robertson for advice on statistics, and S. Morin for helpful edits.

