## Supplemental Material for "Heat dissipation capacity influences reproductive performance in an aerial insectivore"

### Estimated percentage of trimmed surface area

To approximate the proportional amount of surface area that we exposed of the ventral plumage of female swallows, we divided the estimated surface area of the trimmed region ($4.63cm^2$) by the total body surface area of the external plumage ($66.7cm^2$) (4.63 / 66.7 = 6.9%). We used the total surface area of the external plumage of a similarly sized species, the dark-eyed junco (*Junco hyemalis*) (mass = 19.4g, total surface area of exposed plumage = $66.7cm^2$, Walsberg and King 1976) as our estimate for tree swallows (*sensu* Nord et al. 2018). Tree swallow mass during the breeding season ranges from 17 – 22g (Winkler et al. 2011).

### Remote monitoring of activity (reader set-up)

To record parental activity, we used two different radio frequency identification (RFID) readers that received data from the two types of PIT tags (non-temperature sensitive and temperature-sensitive). Non-temperature sensitive tags were read by Generation 2 RFID readers (Cellular Tracking Technology, Pennsylvania, USA) (Bonter and Bridge 2011), while temperature-sensitive tags were read by Biomark HPR Plus readers (Biomark, Idaho, USA). Generation 2 readers (n=50) were placed under each nest box, and were connected to an antenna (125 KHz, #AN0101, QKits, Ontario, Canada) mounted around the entrance hole of the nest box. In this way, individuals were "logged" as they entered and exited the box. We set the 'delay interval', i.e., the length of time that determines how often the same tag can be logged consecutively, to 5s, to avoid generating a large number of reads when a bird sat at the antenna for an extended period of time. Readers were programmed to turn on and off at 0400 and 2200h respectively. We set up antennas at least 24h prior to capturing females for the first time. Biomark HPR plus readers (n=3) were connected to loop antennas (17.5cm), which were positioned so that they encircled the nest box entrance. We cycled the Biomark readers among nests daily so that each nest received a reader for approximately 24h three times throughout the breeding period (early, middle, and late-stage provisioning). Early, middle and late-stage provisioning were defined as days 2-5, 6-9, and 10-14, post-hatch respectively.

### Nestling mass measurements

In 2017, we weighed nestlings on a digital scale at day 0 (Acculab, ± 0.001g) and on a Pesola spring scale (± 0.025g) from day 3 to day 12 post-hatch. In 2018, we weighed nestlings on a digital portable scale (Smart Weigh Digital Pro Pocket, ± 0.01g) from day 0 to day 12. It would be highly unlikely if the use of different scales affected our results, because we had balanced sample sizes between treatments, within years ($Control_{2017}$ = 12, $Trimmed_{2017}$ = 9, $Control_{2018}$ = 18, $Trimmed_{2018}$ = 19) and so this would only influence the estimated effect of 'year' on nestling growth rate. Nevertheless, we weighed a random subset of *adult* birds with both the Pesola and the Smart Weigh scale and found that the mean of the weights was significantly different between scale types (mean of differences = -0.094, $t$= -2.84, 95% CI [-0.16, -0.02], $P$ = 0.012). Despite this statistical difference, a difference of ~0.1g is one small source of variation that could be captured within 'year' and seems likely would have minimal biological meaning relative to the other sources of variation (e.g., weather, date of measurements).

### Nestling growth rates (model selection)

We included 'maternal identity' as a random effect on each parameter (i.e. *A, K, I*) individually and subsequently used an AIC approach to select the optimal random effect structure. We initially ran models with random effects on multiple parameters (e.g., A and K), but these models did not converge and so we assumed that they were over-parameterized and did not consider them further.

**Table S1** Model selection results for determining the optimal random effect structure in the nestling growth rates model. We assigned a random intercept on either the asymptote, inflection point, or growth rate constant. The variance explained by the random intercept was greatest in the model where the random effect was on the asymptote.

| **Random effect structure** | **Delta AIC** | **AIC** | **Log (L)** | **Number of parameters** |
|---|---|---|---|---|
| Asymptote *(A)* | 0.00 | 926.33 | -455.17 | 8 |
| Inflection point *(I)* | 73.98 | 1000.31 | -492.16 | 8 |
| Growth rate constant *(K)* | 106.82 | 1033.15 | -508.57 | 8 |

Tree Swallow (*Tachycineta bicolor*), version 2.0. In The Birds of North America (A. F. Poole, Editor). Cornell Lab of Ornithology, Ithaca, NY, USA. https://doi-org.cat1.lib.trentu.ca/10.2173/bna.11

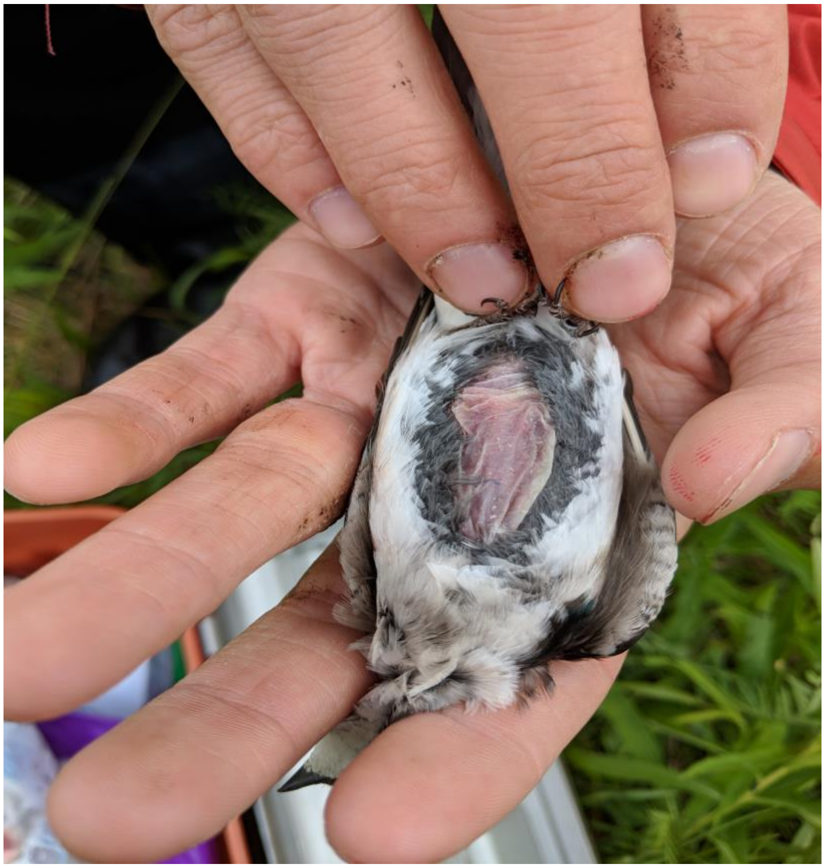

**Figure S1** An example of a female swallow from the trimming treatment. The median amount of ventral region exposed was 3.1mm (length) and 1.9mm (width).
